# Rapid appearance of negative emotion during oral fentanyl self-administration in male and female rats

**DOI:** 10.1101/2023.04.27.538613

**Authors:** Kevin R. Coffey, William Nickelson, Aliyah J. Dawkins, John F. Neumaier

**Affiliations:** Mental Illness Research, Education and Clinical Center, Puget Sound VA Health Care System, 660 S Columbian Way, Seattle, WA 98108; Department of Psychiatry & Behavioral Sciences, University of Washington School of Medicine, Seattle, WA, 98104, USA; Department of Pharmacology, University of Washington School of Medicine, Seattle, WA, 98104, USA

## Abstract

Opioid use disorder has become an epidemic in the United States, fueled by the widespread availability of fentanyl, which produces rapid and intense euphoria followed by severe withdrawal and emotional distress. We developed a new preclinical model of fentanyl seeking in outbred male and female rats using volitional oral self-administration that can be readily applied in labs without intravascular access. Using a traditional two lever operant procedure, rats learned to take oral fentanyl vigorously, escalated intake across sessions, and readily reinstated responding to conditioned cues after extinction. Oral self-administration also revealed individual and sex differences that are essential to studying substance use risk propensity. During a behavioral economics task, rats displayed inelastic demand curves and maintained stable intake across a wide range of fentanyl concentrations. Oral SA was also neatly patterned, with distinct “ loading” and “ maintenance” phases of responding within each session. Using our software DeepSqueak, we analyzed thousands of ultrasonic vocalizations (USVs), which are innate expressions of current emotional state in rats. Rats produced 50 kHz USVs during loading then shifted quickly to 22 kHz calls despite ongoing maintenance oral fentanyl taking, reflecting a transition to negative reinforcement. Using fiber photometry, we found that the lateral habenula differentially processed drug-cues and drug-consumption depending on affective state, with potentiated modulation by drug cues and consumption during the negative affective maintenance phase. Together, these results indicate a rapid progression from positive to negative reinforcement occurs even within an active drug taking session, revealing a within-session opponent process.

**Significance Statement:** The United States opioid epidemic is defined by rampant and treatment resistant fentanyl use. Better understanding of neural substrates underlying this phenomenon is essential to slowing the opioid crisis. Intravenous and vapor self-administration (SA) are the standard models for studying fentanyl use in rodents, however they many carry pragmatic downsides. Here, we used a novel oral fentanyl self-administration model that provides key translational and technical benefits and can be readily applied in other labs to study the neurobiology of fentanyl SA. This method captured individual and sex differences necessary for studying substance use risk propensity and uncovered a rapid shift in affective state in rats, suggesting and shift from positive to negative reinforcement within each fentanyl taking session.

## Introduction

Opioid use disorder is an epidemic in the United States, where fentanyl has caused a dramatic spike in overdose deaths (>70,000/year (CDC, 2022)). Fentanyl’ s low cost, ready availability, and extreme potency contribute to its profound impacts and difficult treatment (Pergolizzi et al., 2021; Reuter et al., 2021; Stanley, 2014). It produces powerful but fleeting euphoria, while withdrawal causes severe physical and emotional turmoil. In abstinence, strongly conditioned drug-cues and hypersensitivity to emotional distress motivate persistent relapse (Koob, 2020). Careful preclinical modeling of volitional fentanyl intake is essential to understanding both the adaptations in neurobiology caused by chronic drug exposure and the individual differences in neurobiology that could predict addiction susceptibility.

Currently, intravenous (IV) self-administration (SA) is considered the gold standard model for studying volitional drug intake in rodents, and vapor SA is becoming increasingly popular (Moussawi et al., 2020; Vendruscolo et al., 2018). However, IV and vapor SA come with pragmatic downsides such as invasive surgeries, catheter maintenance, and dangers associated with aerosolized fentanyl. An oral fentanyl SA approach could circumvent these issues while providing additional translational benefits. Half of fentanyl users swallow or insufflate (Jiang et al., 2021), and individuals who abuse prescription opioids tend to chew their pills to accelerate absorption (Gasior et al., 2016; Jiang et al., 2021). Rodents drinking small boluses of dissolved fentanyl faithfully model these routes of administration, as fentanyl citrate is readily absorbed into the oral mucosa and rapidly diffuses across the blood brain barrier (Naji & Ramsingh, 2023). Oral fentanyl SA could also be valuable for studying individual and sex differences. Women are more likely to consume opioids orally (Back et al., 2011), and females rodents consume increased quantities of oral oxycodone (Phillips et al., 2020).

Here, we endeavored to develop a robust and convenient oral fentanyl SA model that is useful for studying volitional fentanyl intake in males and female rats. Animals in our model showed escalating intake, strong association of conditioned cues, and reinstatement, along with key individual and sex differences. They also displayed inelastic demand curves, maintaining stable intake across a wide range of doses. Oral fentanyl SA produced patterned SA similar to IV cocaine, with distinct “ loading” and “ maintenance” phases (Coffey et al., 2015; Root et al., 2011). During loading, animals responded rapidly to increase their drug level, then switched to slower maintenance responding to hold a stable drug-level. Using our recently released software DeepSqueak v3 (Coffey et al., 2019), we captured and categorized thousands of ultrasonic vocalizations (USVs) during self-administration, extinction, and reinstatement. USVs are innate expressions of current emotional state in rats (Burgdorf et al., 2020; Knutson et al., 2002) and were used to characterize the relationship between drug taking phase and affective state. Positive-affective 50 kHz USVs were produced during the loading phase, while negative-affective 22 kHz USVs were produced during maintenance, suggesting animals shift from positive to negative reinforcement within individual sessions (Barker et al., 2014).

Oral SA is ideal for pairing with complex neurophysiological techniques. Here, we recorded calcium activity in the lateral habenula (LHb) using fiber photometry during oral fentanyl SA. The LHb, a region known to process both aversion and the motivational value of reward related cues (Bromberg-Martin & Hikosaka, 2011; Hikosaka, 2010), responded to drug-cues and consumption differentially depending on the phase of SA. Changes in LHb activity were potentiated during the negative affective maintenance phase compared to loading. Together, the USVs and LHb activity reveal a rapid shift from positive reinforcement during the loading phase to a negative reinforcement during the maintenance phase, occurring much earlier than generally thought (Dao et al., 2021; Koob, 2020). Furthermore, oral fentanyl SA provides substantial translational and technical benefits, enables the study of neurobiological, motivational, and emotional aspects of fentanyl use, and captures valuable individual and sex differences related to substance use risk propensity. This model also provides a level of convenience that could democratize access to fentanyl SA procedures and accelerate neurobiological discoveries.

## Results

### Oral Fentanyl Self-Administration is a Robust & Convenient Model in Male & Female Rats

Female (n=12) and male (n=10) rats were trained to self-administered oral fentanyl 5 days a week, 3 hours a day, for 15 days, followed by 7 days of extinction training and a single cued-reinstatement session (Figure 1A). Both male and female rats escalated drug seeking across training (F(1,308)=96.325, p<.001), rapidly decreased responding during extinction (F(1,132)=63.20, p<.001), and increased responding again during cued reinstatement (F(1,22)=41.47, p<.001). Females also had higher rates of extinction responding (F(1,138)=6.00, p<.05) and reinstated to fentanyl cues at nearly double the rate of males (F(1,22)=5.79, p<.05; Figure 1B). Inactive lever pressing remained low throughout oral SA (Figure 1C). The total number of rewards (F(1,308)=148.77, p<.001; Figure 1D) and total intake of fentanyl (F(1,308)=231.41, p<.001; Figure 1E) both significantly increased across training for male and female rats. Male rats made significantly more head entries throughout training (F(1,40)=13.37, p<.001), though both male and female rats rapidly reduced head entries during extinction (F(1,132)=49.65, p<.001) and increased head entries during reinstatement (F(1,22)=20.76, p<.001; Figure 1F). Finally, rats developed a strong association between the cued lever press and reward consumption, reducing their latency from lever press to head entry over sessions (F(1,308)=63.72, p<.001; Figure 1G). Animals showed increased latency variability during extinction, and significantly reduced latency during reinstatement testing (F(1,22)=8.36, p<.01; Figure 1G).

**Figure 1.**
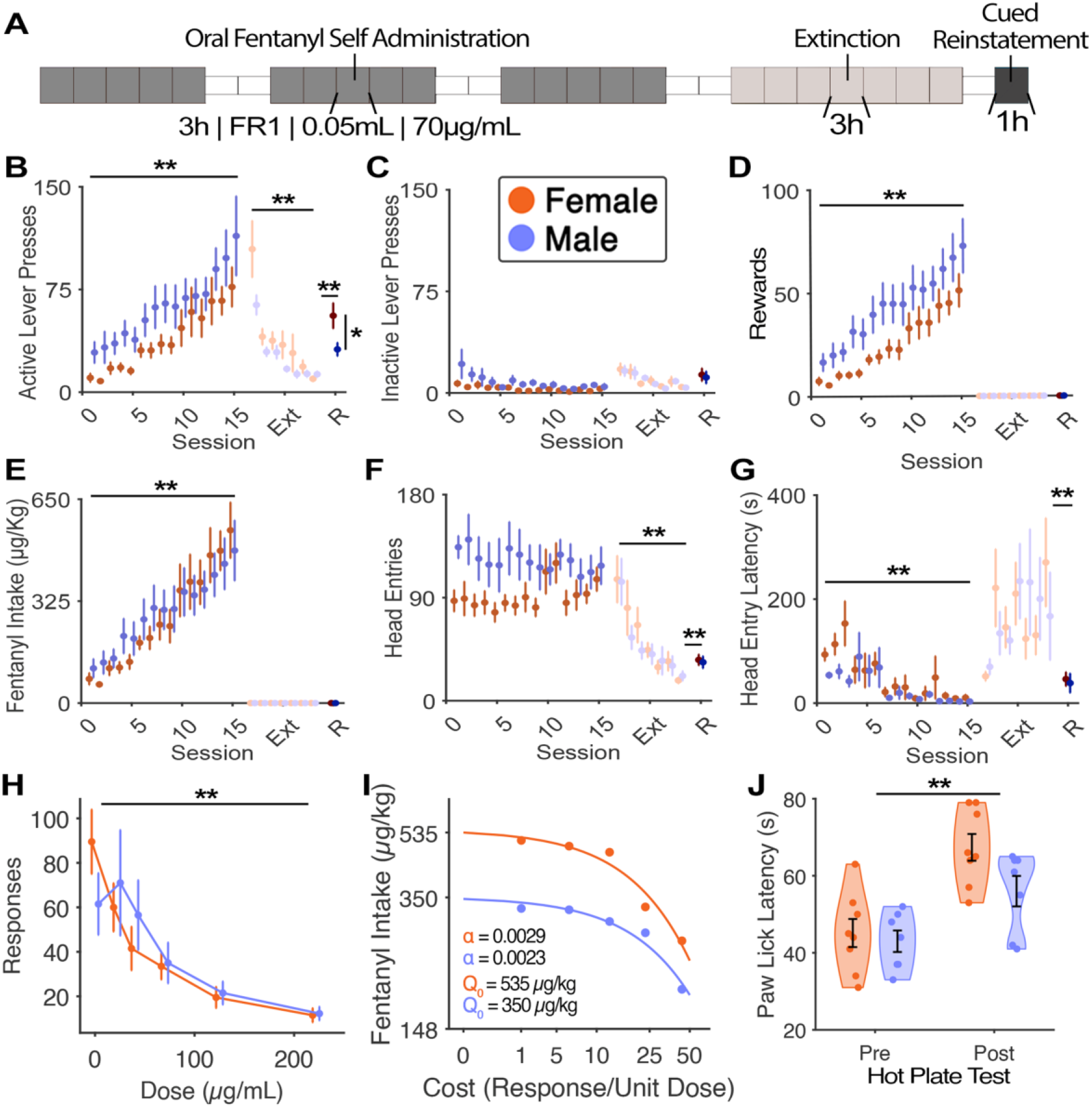
Oral Fentanyl Self-Administration. | **A)** Timeline of oral fentanyl self-administration (SA), extinction, and cued-reinstatement testing. Self-administration behavior in male (n=10) and female (n=12) rats including **B)** active lever presses, **C)** inactive lever presses, **D)** total rewards earned, **E)** total fentanyl consumed, **F)** total head entries, **G)** and head entry latency. A separate cohort of male (n=8) and female (n=8) rats underwent between-sessions behavioral economics and hot-plate procedures. **H)** Total responses at varying fentanyl concentrations. **I)** Demand curves highlighting intake at zero cost (Q0) and demand elasticity (α). **J)** Paw lick latency before and after one hour of oral SA at 70 μg/mL fentanyl demonstrates a decreased sensitivity to pain after fentanyl taking in both sexes. In all panels, orange data reflects females and purple data reflects males. ^*\**^*p<0*.*05*, ^*\*\**^*p<0*.*01*.

### Oral Fentanyl SA Produces Antinociception and Inelastic Demand Curves

To further validate our oral SA model, a separate cohort of female (n=8) and male (n=8) rats were tested on a modified between-sessions thresholding procedure. During the third week of SA, the dose of fentanyl decreased between each session (Oleson & Roberts, 2009). The dose started at 222 μg/mL and decreased daily on a quarter-log scale (222, 125, 70, 40, 22, 0 μg/mL). As the dose of fentanyl decreased, rats increased their rate of responding (F(1,80)=42.46,p<.001; Figure 1H). For females, estimated fentanyl intake at zero-cost (Q0) was 535 μg/kg, while for males Q0 was slightly lower at 350 μg/kg (Figure 1I). Demand was relatively inelastic in both sexes, and fentanyl intake remained stable in males and females above 70 μg/mL (Figure 1I), though females displayed slightly more demand elasticity than males (female α=0.0029, male α=0.0023). Finally, these animals were tested to determine if oral fentanyl SA was antinociceptive. After 1 hour of 70 μg/mL SA, when drug levels have stabilized, males and females showed a significant increase in paw lick latency during a hot-plate test (F(1,15)=31.98, p<.001; Figure 1J).

### Male and Female Rats Show Distinct Fentanyl Use Risk Trajectories

A descriptive analysis of individual differences in multiple behaviors was performed to ascertain the underlying behavioral distribution of fentanyl use risk severity. Both male and female rats were roughly normally distributed across multiple fentanyl use behaviors. Kolmogorov-Smirnov tests against the standard normal distribution failed to reject the null hypothesis for fentanyl intake (p=.55; Figure 2A), fentanyl seeking (p=.69; Figure 2B), cue association (p=.32; Figure 2C), escalation of intake (p=.81; Figure 2D), extinction responding (p=.11, Figure 2E), and relapse (p=.66; Figure 2F). These behaviors are also relatively cross-correlated. Spearman’ s rank-order correlation revealed that total intake was most correlated with other fentanyl use behaviors, while persistent responding during extinction was most correlated with cued-reinstatement responding (Figure 2G). By combining these different behaviors, we can assign each animal a composite fentanyl use severity score, which also appeared to be normally distributed (p=.87; Figure 2H). Using a principal component analysis of the combined behavioral characteristics revealed that “ high-severity” male and female rats had different underlying behavioral trajectories, with high-severity male rats prone to high seeking, escalation, and total intake, while high-severity females were better defined by strong cue-association, persistent extinction responding, and higher relapse rates.

**Figure 2.**
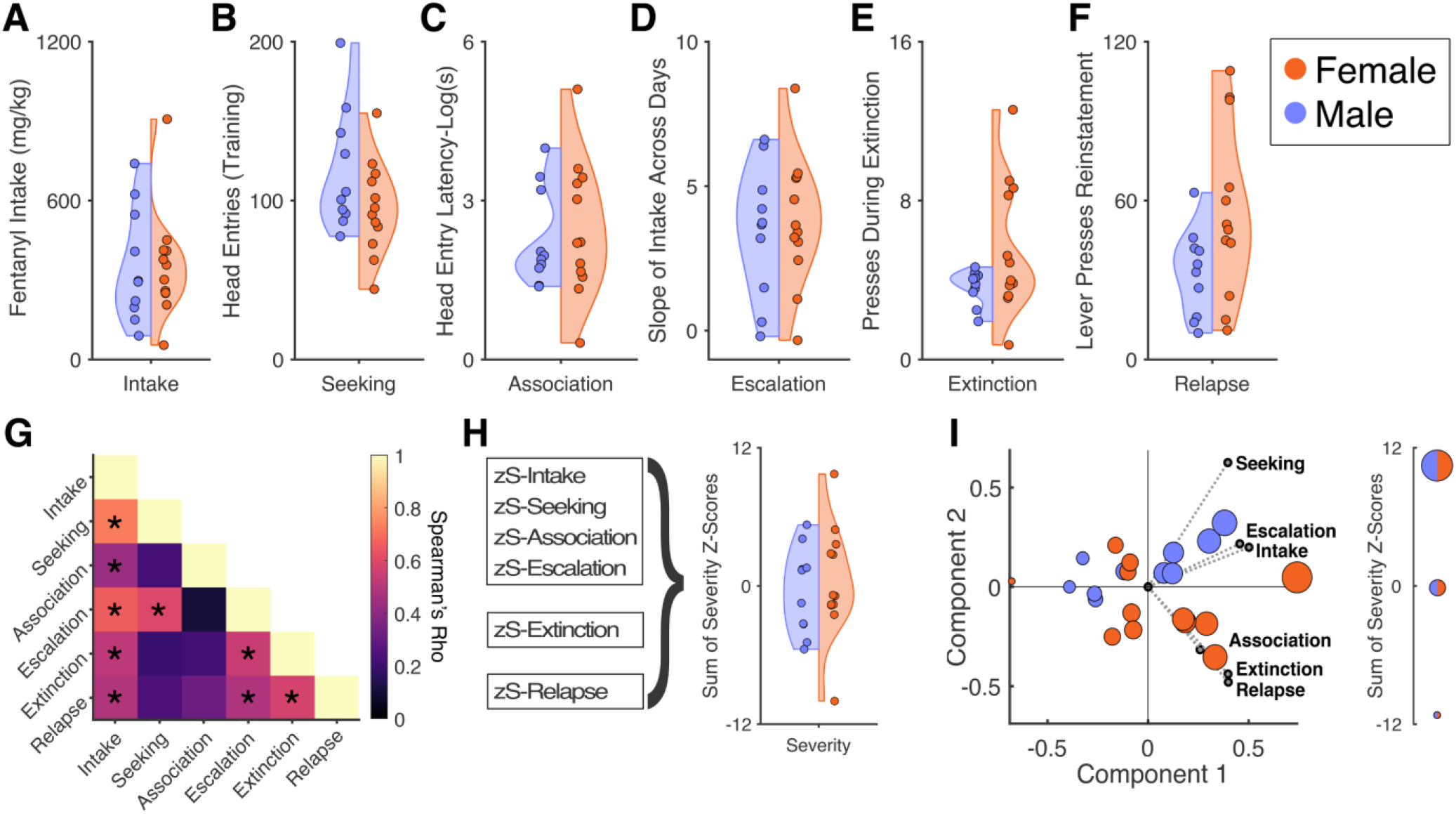
Individual differences In Self-Administration Behavior and Fentanyl Use Severity |. Behavioral distributions for different componenets of fentanyl use severity, including **A)** fentanyl intake, **B)** fentanyl seeking (defined as total head entries), **C)** cue-association (defined as head entry latency), **D)** escalation (defined as slope of total intake), **E)** persistence in extinction (defined as total presses during extinction), and **F)** relapse (defined as total pressed during cued reinstatement). **G)** Cross-correlation of individual risk severity measures. H) Calculation of individual fentanyl use severity scores. I) Principal component “ Bi-Plot” analysis showing high-risk male and female rats in separate quadrants associated with different components of fentanyl use severity. In all panels, orange data reflects females and purple data reflects males. ^*\**^*p<0*.*05*.

### 50 and 20 kHz Ultrasonic Vocalizations Show Distinct Temporal Structure During Oral Fentanyl SA

Rats make a wide variety of ultrasonic vocalizations during oral fentanyl self-administration. Using unsupervised clustering on a weighted combination of extracted contours (Coffey et al., 2019) and latent features defined via variational auto encoder (Goffinet et al., 2021), USVs were automatically clustered into 12 categories (Figure 3A). As has been shown previously, calls in the 38-90 kHz range (50 kHz) do not cluster into well segregated categories, and instead exist along a continuum in feature space (Goffinet et al., 2021), while calls in the 18-38 kHz range (22 kHz) are well separated from the others (Figure 3B). Still, it can be valuable to categorize 50 kHz calls to determine if they are used differentially in certain contexts. Here, we find that all categories of 50 kHz and 22 kHz USVs are used throughout each phase of SA, including training, extinction, and reinstatement (Figure 3C,D,E). However, the time-course of their use differs throughout the SA phases. During training, all types of 50 kHz USVs are expressed early in the session, beginning just prior to the first infusion, then tapering off by 30 minutes. At ∼30m, animals begin to produce 22 kHz USVs (Figure 3F). During extinction training a similar pattern is observed, but the shift occurs in quicker succession. As no drug or cues are presented in extinction, the animals may become frustrated more quickly (Figure 3G). During reinstatement we see a similar pattern to training, perhaps because the return of the fentanyl associated cues extends the production of 50 kHz USVs (Figure 3H).

**Figure 3.**
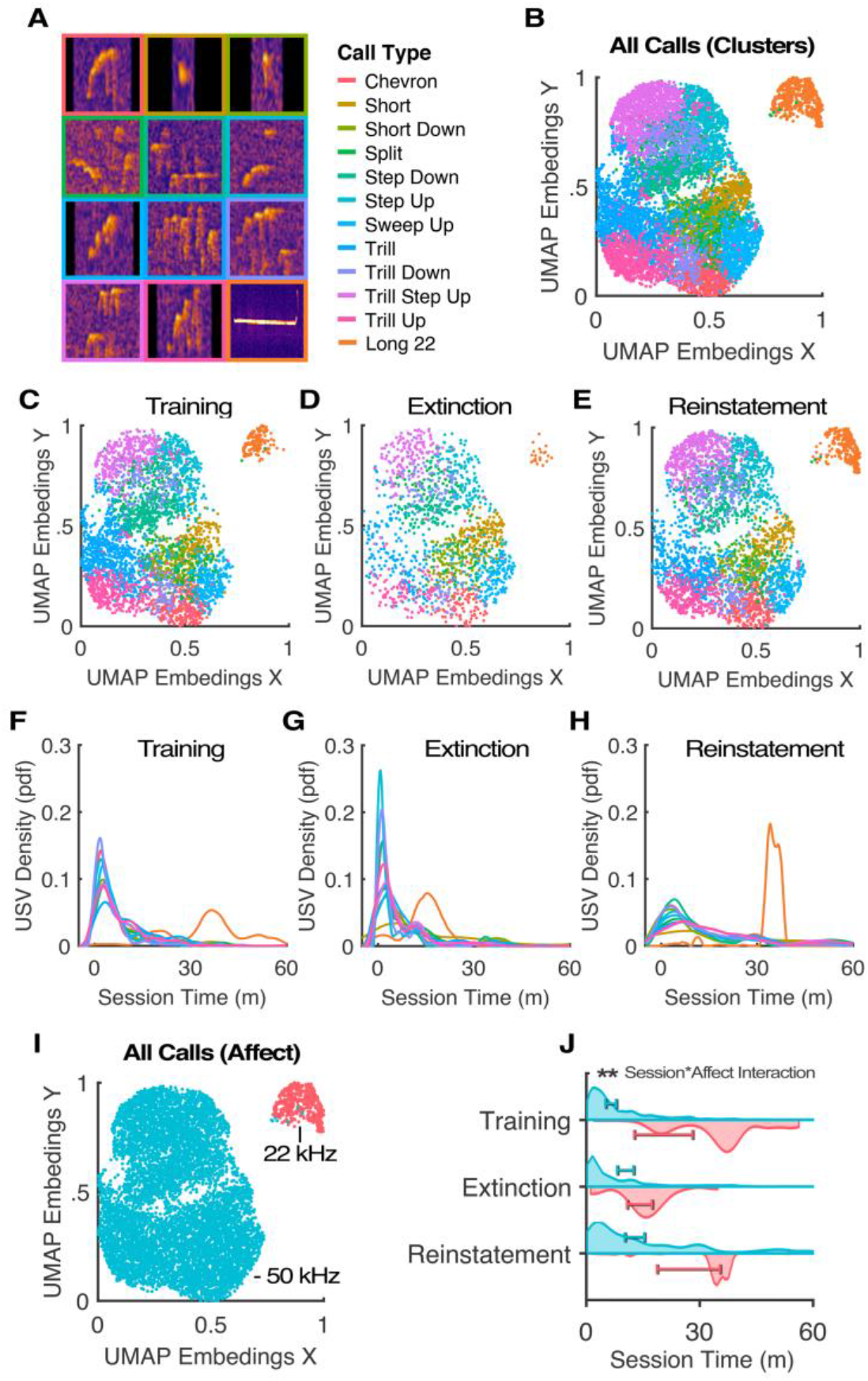
USV Evidence for an Affective Opponent Process during Fentanyl SA |. **A)** Automatic classification of 12 USV categories based on variational autoencoder embeddings and contour parameterization. **B)** Projection of USV categories in 2D space using UMAP to reduce latent feature space. All categories of USVs are used across each phase of fentanyl self-administration, including **C)** training, **D)** extinction, and **E)** reinstatement testing. However, the time course of long 22 kHz USVs differs from all other USV categories during **F)** training, **G)** extinction, and **H)** reinstatement. **I)** Simplified categorization of USVs into positive and negative affective calls based on well-established frequency criteria. **J)** Positive affective USVs are used early in each session during drug loading, while negative affective calls are produced later, during maintenance. In panels I and J, red data reflects ∼22 kHz and blue data reflects ∼50 kHz calls. ^*\*\**^*p<0*.*01*

Since all the types of 50 kHz USVs followed a similar pattern throughout SA, we chose to collapse them into their affective representations, whereby calls >38kHz are defined as “ positive affective” and calls <38kHz were defined as “ negative affective” (Burgdorf et al., 2011; Portfors, 2007). These categories are well defined functionally and in feature space (Figure 3I), and the timing of their production could be compared across session type. Here we found a significant interaction of session type and affect category on call time (F(1,67)=11.24, p<0.01), whereby positive affective USVs are produced early in the session, followed by a rapid shift to negative affective USVs.

### The “ Loading” & “ Maintenance” Phases of Oral Fentanyl SA Reflect a Transition in Affective State Consistent with Opponent Process Theory

Oral fentanyl self-administration appears composed of two distinct phases, a “ loading” phase where animals rapidly consume drug to reach a particular internal drug level, followed by a “ maintenance” phase where that drug level is maintained with slower, evenly spaced consumption. This pattern can be seen in the estimated brain fentanyl concentration of individual animals (Figure 4A) along with their corresponding cumulative response distributions (Figure 4B). If we combine this data for all animals, it is evident that, while the maintenance drug level increases across sessions, the pattern remains the same (Figure 4C). Animals accomplish this by increasing the number and rate of operant responses during both the loading phase and the maintenance phases in later sessions as training proceeds. Because of the exponential nature of pharmacokinetics, maintaining a higher drug level requires an increase in response rate. While the shift from loading to maintenance is defined by the shift in response rate, peak drug level occurs sometime later due to continued absorption and transfer across the blood brain barrier (Figure 4E). This transition from loading to maintenance coincides with a shift from positive affective to negative affective vocalizations (Figure 4E,F). Interestingly, the peak of negative affective vocalization production occurs concurrent with the peak of estimated brain fentanyl concentration (Figure 4E,F), providing corroborating evidence that our coefficients for fentanyl pharmacokinetics are accurate.

**Figure 4.**
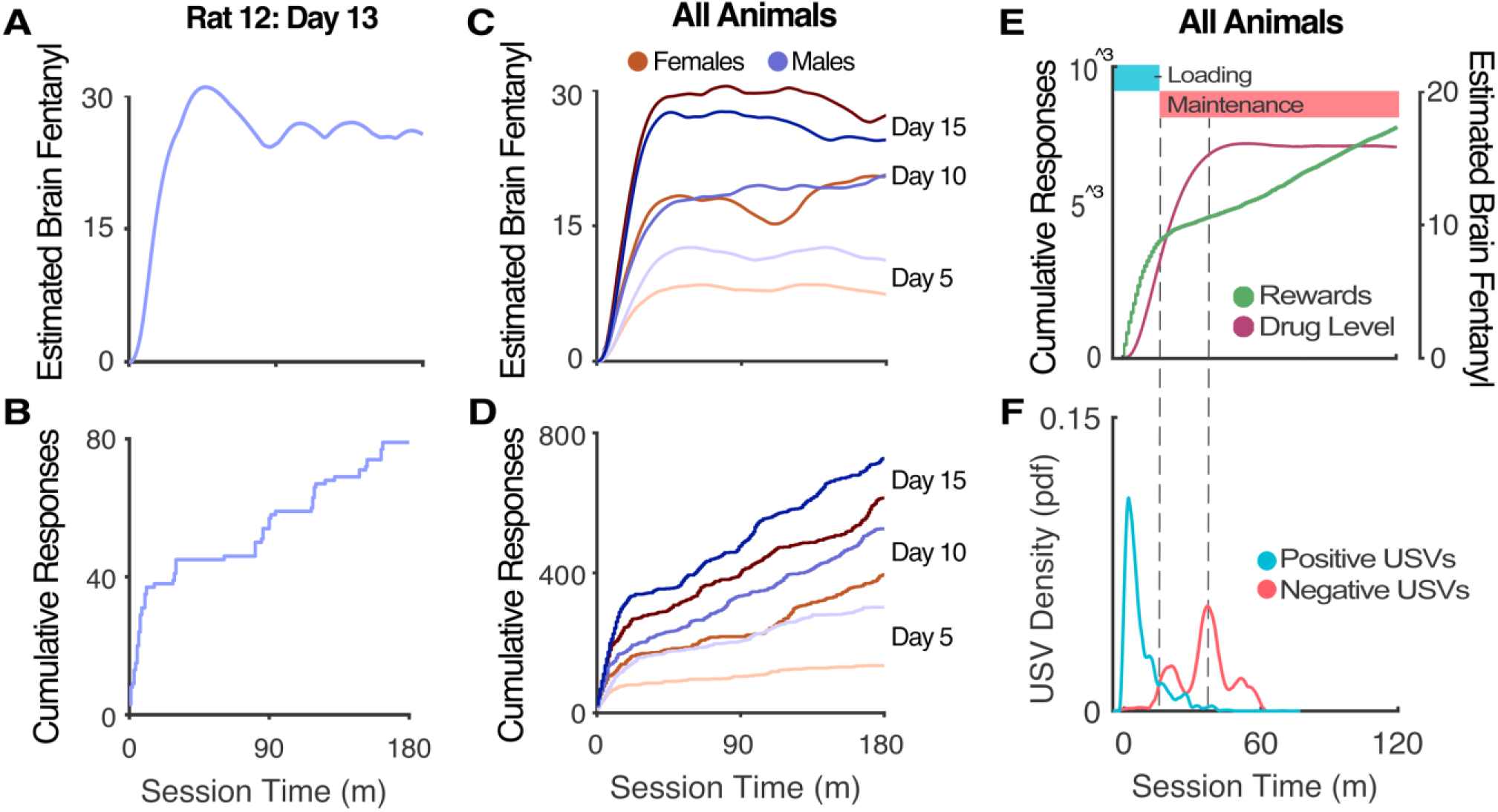
Oral Fentanyl SA Has Distinct Positive-Affective Loading and Negative-Affective Maintenance Phases |. **A)** Individual example of estimated brain-fentanyl concentration and **B)** corresponding cumulative response pattern, together showing distinct loading and maintenance phases. **C)** Average estimated brain-fentanyl concentration for all males and females on days 5, 10, and 15. **D)** Corresponding cumulative response pattern for all animals on days 5, 10, and 15. **E)** Average estimated brain-fentanyl concentration for all animals on all training days and corresponding cumulative response pattern. Loading and maintenance are defined by the elbow of the cumulative response curve, where the rate of responding clearly shifts. **F)** USV probability density function showing all USVs from all animals. 50 kHz USVs occur during loading, then rapidly shift to 22 kHz USVs which peak concurrent with max drug level. Orange data reflects females and purple data reflects males, while blue data reflects ∼50 kHz calls (loading) and red data reflects ∼22 kHz calls (maintenance). Green data reflects fentanyl reward deliveries, and pink data reflects estimated brain fentanyl concentration.

### LHb Differentially Processes Fentanyl Cues and Rewards Dependent on Self-Administration Phase

As the lateral habenula is known to process information about rewards, cues, and aversive states (Bromberg-Martin & Hikosaka, 2011; Hikosaka, 2010; Lecca et al., 2014), we used fiber photometry to examine whether the LHb is modulated by rewards and cues differently during the positive affective “ loading” phase and the negative affective “ maintenance” phase of SA. We found that LHb activity is significantly increased by the novel conditioned stimulus (CS) early in training, and that this increase was greater and more sustained during the maintenance phase (Figure 5E). Over weeks of training, this cue-induced activity diminishes, and is only present during the maintenance phase in weeks 2 and 3 (Figure 5F,G). Intriguingly, the LHb did not show any response to the CS during reinstatement testing (Figure 5H). Early in training, when animals were still forming the cue-reward association, LHb activity fell during reward consumption (rewarded head entry; Figure 5I). This decrease was only present during the maintenance phase in week 2 (Figure 5J) and was absent in week 3 (Figure 5K).

**Figure 5.**
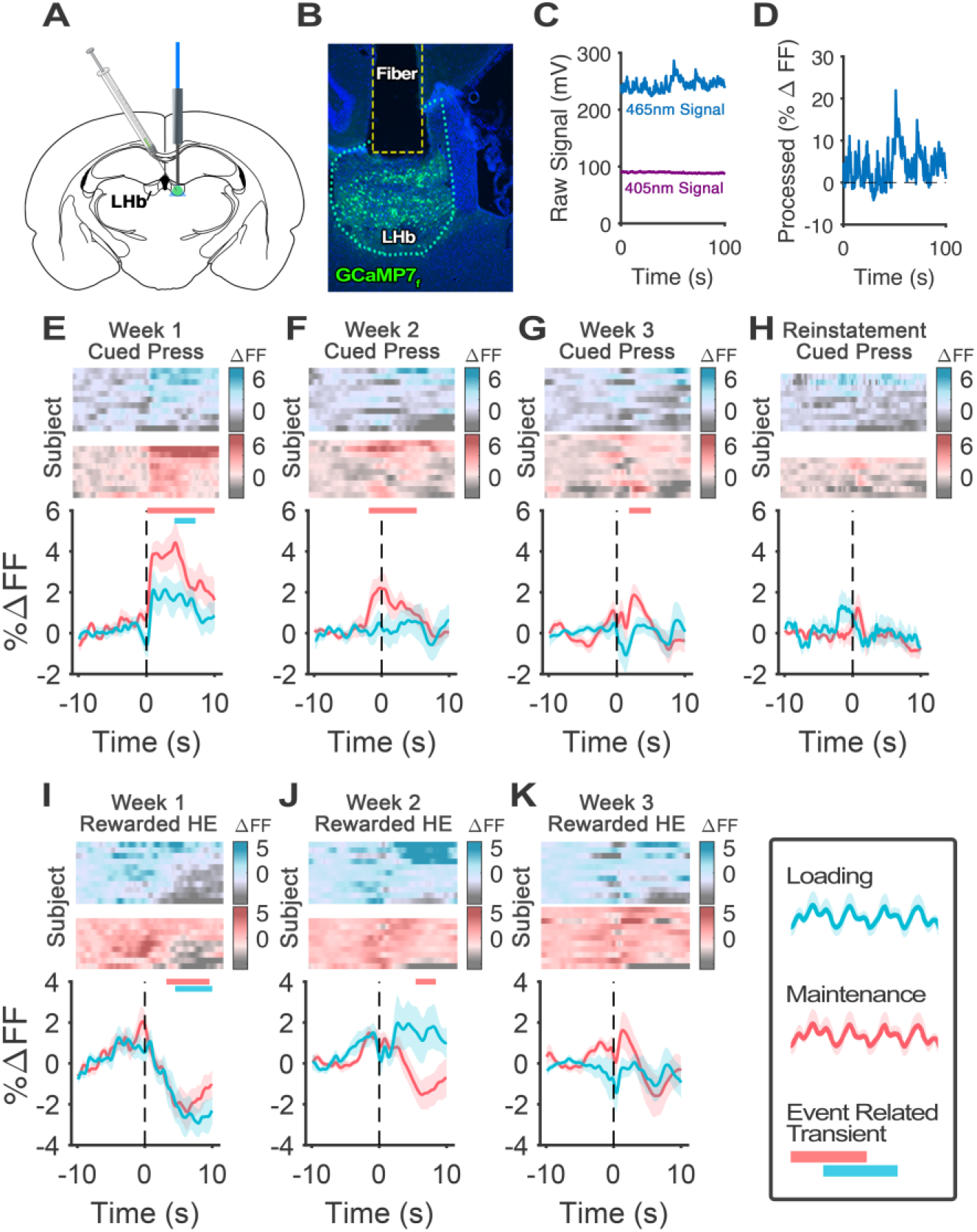
LHb Differentially Processes Fentanyl Cues and Consumption Dependent on Self-Administration Phase |. **A)** For fiber photometry, rats received pGP-AAV1-syn-jGCaMP7f-WPRE into the right LHb, followed by a borosilicate optical fiber. **B)** Histological verification of injection targeting and fiber placement. **C)** Example raw trace from the isosbestic control 405 nm LED and calcium signal generating 465 nm LED. **D)** Processed fluorescence signal (%ΔFF) was used to compare across animals and conditions. **E)** LHb activity increased to the cued lever press early in training, but this signal diminishes over training, and is only present during maintenance during **F)** week 2 and **G)** week 3 and is completely absent during **H)** reinstatement testing. **I)** LHb activity decreases during fentanyl consumption (rewarded head entry) early in training, but this signal diminishes over training, and is only present during maintenance during **J)** week 2 and is completely absent during **K)** week 3. Blue data reflects LHb activity from loading and red data reflects LHb activity from maintenance. *Bootstrap CIα for ERTs = 0*.*05*.

## Discussion

Fentanyl use disorder in humans is a complex behavioral phenomenon with multiple components, including positive and negative reinforcement, emotional dysregulation, escalating intake, and persistent relapse risk. To glean new insights into opioid neurobiology and provide new treatments for fentanyl use disorder, researchers rely on preclinical models, which must faithfully capture these defining components to be translationally relevant. The current ‘ gold-standard’ for preclinical models is long-access intravenous (IV) self-administration (SA), extinction, and reinstatement testing. This model faithfully captures escalation, the affective opponent process, and relapse behaviors. While there are strengths to IV models, they also present several disadvantages, including additional surgery, catheter patency issues, and single housing stress. Here, we present our newly developed oral fentanyl SA model, which dramatically simplifies SA procedures, while retaining the relevant translational features.

### Oral Fentanyl Self-Administration Produces Clinically Relevant Escalation of Intake and Reinstatement

Oral fentanyl SA occurs in two distinct phases: a “ loading” phase characterized by rapidly elevating brain-drug concentrations, followed by a prolonged “ maintenance” phase characterized by equally spaced responses that maintain a stable brain-drug concentration. In our model, escalation of intake occurs as animals learn to load faster and reach higher levels of maintained intake, leading to significantly increased levels of operant responding and total intake in both males and female rats. By week 3, rats are consuming ∼450 μg/kg in 3h, well in excess of intake in 6h IV SA sessions (150-375 μg/kg; (Dao et al., 2021; Hammerslag et al., 2021; Malone et al., 2021)). This is consistent with the generally observed 50% bioavailability of oral fentanyl citrate (Mystakidou et al., 2006; Naji & Ramsingh, 2023). Operant responding in our paradigm appears specific for fentanyl, and not the water vehicle, as rats maintain stable intake across a wide range of doses (and water volumes) by modulating response rate. This suggests that animals’ primary goal while self-administering is to maintain a particular brain-fentanyl concentration, similar to what has been observed with IV cocaine SA (Oleson & Roberts, 2009; Root et al., 2011). After SA and extinction training, animals undergo a test for reinstatement of seeking. When conditioned cues are returned, rats readily resumed lever pressing even in the absence of reinforcement. Taken together, these data provide strong evidence that oral fentanyl SA is a valid preclinical model for studying escalation of fentanyl intake and cued relapse to seeking in male and female rats (Schuster & Woods, 1968; Stewart et al., 1984).

### Rats Display Individual & Sex Differences in Oral Fentanyl Seld-Administration

Modern addiction neuroscience is rooted in the idea that drugs of abuse hijack and reshape the brain’ s natural reward system (Bliss et al., 2003; Spanagel & Weiss, 1999; Volkow & Li, 2005). It is interesting to note however, that while these neural adaptations occur across whole groups of drug users, only a small subset of those who try drugs will go on to develop problematic use (Schlag, 2020). Indeed, there is evidence from genetically-heterogenous rodent studies and human self-report studies that individual variability accounts for substantial differences in patterns of drug use (Bagley et al., 2022; Schlag, 2020). The development of preclinical models that capture these individual differences is essential to the study of neurobiological correlates of addiction risk and susceptibility.

Male and female rats in our fentanyl SA model show substantial individual and sex difference that can be used for assessment of fentanyl use risk severity. Rats showed wide but relatively normal distributions of fentanyl intake, seeking, cue association, escalation, and relapse. Though these behaviors are somewhat cross-correlated, they are distinct enough to be leveraged for principal component analysis and used to extract the features that contribute to overall fentanyl use risk severity. In our analysis, the males with the highest risk severity scores were primarily driven by heightened escalation, total intake, and fentanyl seeking (head entries), while the highest risk severity females were defined by increased cue association (head entry latency), extinction responding, and reinstatement responding. We also found that individual rats displayed demand curve inelasticity, which provided additional insight into each animal’ s motivation to take fentanyl. In future studies, these behavioral differences will be valuable for identifying underlying differences in genetic or neuronal processing that predict drug abuse liability and could improve addiction prevention or individualize addiction treatment based on deeper understanding of the brain processes accounting for different patterns of misuse.

### Oral Fentanyl Self-Administration Produces Within-Session Affective Opponent Process

The affective opponent process theory of addiction postulates that addiction is driven by two opposing processes: the initial pleasurable effects of drugs (process-A), and a subsequent negative affective state that arises as a result of repeated drug use (process-B) (Solomon & Corbit, 1974). This leads to a cycle of addiction, where individuals seek drugs to alleviate negative affective states, despite the negative consequences of drug use. This theory is well supported in the literature (Heilig et al., 2010; Kassel et al., 2007; Koob, 2017; Wheeler et al., 2008), and has been shown to occur during IV fentanyl SA in rats (Dao et al., 2021). Both sensitization and tolerance can shape these dueling features over time, but it was believed that some cessation of drug intake was a critical factor in process-B. Here, we provide evidence that the affective opponent process can be detected even when drug taking continues and estimated brain concentrations fluctuate only slightly.

Animal’ s maintained drug level during SA has been framed as “ satiety” and was presumed to be the drug level at which animals were satisfied, or comfortable (Root et al., 2011; Zimmer et al., 2011). However, evidence from ultrasonic vocalizations produced during cocaine self-administrations studies painted a different picture. Animals primarily produced 50 kHz “ positive” affective USVs during the initial loading phase, and 22 kHz “ negative” affective USVs in satiety (Barker et al., 2014). Here we replicate and extend these results to oral fentanyl SA. In this study, rats produced a myriad of 50 kHz vocalizations during the loading phase, but when responding slows during maintenance they produced almost exclusively negative affect-associated 22kHz vocalizations. The transition from 50 to 22 kHz vocalizations occurs precisely at the time when response rate shifts from loading to maintenance, and the peak in 22 kHz USVs occurs a short time later, when drug level also peaks. Take together, these data suggest that the maintenance phase is defined by a narrow range of drug-level, above or below which animals experience negative affect, motivating behavior to return to that range. These features make oral fentanyl SA a compelling model for studying emotional motivation and negative reinforcement in opioid use disorder.

### LHb Differentially Processes Fentanyl Cues and Rewards Dependent on Affective State

The lateral habenula is a small but essential node in limbic to midbrain circuitry (Bernard & Veh, 2012) that is known to process the motivational and emotional aspects of substance use, withdrawal, and relapse (Clerke et al., 2021, 2021; Lecca et al., 2014; Wang et al., 2017). To test our model’ s usefulness for neurophysiological studies, we recorded calcium signals in the LHb using fiber photometry across oral fentanyl SA, extinction, and reinstatement. We found that LHb activity was primarily modulated by cues and rewards early in training when animals were still forming the cue-fentanyl association. The LHb was strongly activated by novel cues during week 1, and significantly more so during the maintenance phase when animals were in a negative affective state. This suggests that increased LHb activity was not due primarily to acoustic startle, as LHb activity to late session cues was potentiated as compared to the more novel, and ostensibly startling, early session cues. Thus, the negative affective state experienced in maintenance may provide extra salience to the CS that modulates LHb’ s responsivity. This pattern is similarly reflected in the LHb’ s modulation by fentanyl consumption. While the CS increased LHb activity, consumption of the fentanyl reward produced a rapid decrease in LHb activity. This change is similar during loading and maintenance in week 1, but is only present during maintenance in week 2, providing further evidence that the affective transition during maintenance modulates LHb’ s sensitivity to fentanyl cues and consumption. By week 3, the lever press, cue, and consumption are tightly temporally linked, making it difficult to separately analyze these behaviors. Though small, the LHb may play a critical role in the affective opponent processes driving substance use disorder, and a better understanding of its role may lead to new treatments.

### Conclusions and Future Directions

Here, we present a novel oral fentanyl self-administration model aimed at democratizing access to fentanyl SA procedures and accelerating discoveries that improve treatment for fentanyl use disorder. Our model faithfully recapitulates key components of fentanyl use disorder, such as escalation of intake, cue association, relapse, demand inelasticity, and the affective opponent process. It also provides a level of convenience that makes proposing and initiating new studies cheaper, faster, and more efficient compared to IV or vapor SA. We hope that by combining this model with cutting edge neurophysiology and genetics toolkits, we can gain new insights into the neuronal adaptations caused by chronic fentanyl use and the underlying individual differences in neurobiology that predict addiction susceptibility.

## Methods

### Animals

For all experiments, male and female Long-Evan rats (Charles River) weighing 250-450 g were used. Rats were dual-housed in a temperature and humidity-controlled room on a 10-14 dark-light cycle and acclimated to the facility for 1 week before experimental procedures. Food and water were available ad-lib. Experimental procedures were approved by the University of Washington Institutional Animal Care and Use Committee.

### Experimental Design: Oral Fentanyl Self-Administration & Reinstatement Testing

Male (n=10) and female (n=12) rats (Long Evans, Charles River) were trained to self-administer (SA) liquid fentanyl, while a subset of those animals (n=12) was used for fiber photometry. Rats learned to SA fentanyl (70 μg/mL) in H2O (0.05 ml/delivery) for 5 days a week (3 hr/session) over three weeks on an FR1 schedule (Figure 1A). The first 5 days of SA employed a “ saccharin fade” (0.1%, 0.08%, 0.06% 0.04%, 0.02%) where the full dose of fentanyl is present from day 1. Upon active lever press, liquid fentanyl was delivered to a small dish paired with a 10-second presentation of an audiovisual conditioned stimulus (CS) consisting of illumination of a red light above the active lever for 1-second coincident with a 10-second tone (4 kHz) and 10-second extinction of the house light. During the CS presentation a time-out was imposed. SA occurred in standard operant chambers (Med-Associates) with 2 levers. Presses on the second (inactive) lever were recorded with no programed consequences. On day 22, rats began extinction training wherein lever pressing resulted in neither cue presentations nor fentanyl delivery. Extinction sessions (3 hr/session) continued for 7 days for all animals. Twenty-four hours after the final extinction session, animals underwent cued reinstatement testing (1 hr/session) where the compound CS signaled the beginning of the session. Each subsequent active lever press elicited the compound CS but did not result in the delivery of fentanyl.

### Experimental Design: Between Session Thresholding & Hot Plate Testing

A separate cohort of male (n=8) and female (n=8) rats was used to further validate our oral-fentanyl SA model with a hot-plate test for antinociception and a between-sessions-thresholding procedure to explore demand elasticity and dose-intake characteristics (Oleson & Roberts, 2009). The week before SA, rats underwent a hot-plate pre-test to determine baseline nociception. Rats were placed on a precision hotplate (52°C) and latency to lick their rear paws or jump to the chamber rim (12”) was calculated (max time= 90s). The following week, animals were trained to SA liquid fentanyl in the manner described above. On day 8 of SA, animals were removed from the chamber 1h into the SA session and placed on the hotplate for the post-test. On day 11 of SA, animals began the between-sessions-thresholding procedure. For this procedure, the concentration of fentanyl was progressively decreased each day for 6 days following a (¼)Log series (222 μg/mL, 125 μg/mL, 70 μg/mL, 40 μg/mL, 22 μg/mL, 0 μg/mL). Unlike normal SA sessions, animals did not always consume all of the liquid that was delivered, particularly on high concentration days. However, because liquid was delivered via a syringe pump to a deep dish where no spillage occurred, we were able to draw back the remaining liquid at the end of each session to measure total consumption precisely. Demand curves for male and female animals were fitted to raw fentanyl intake data using the following equation:

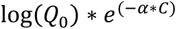

Here, Q0 and α are model coefficients and C is the independent variable. C represents the unit cost of fentanyl, where 1 lever press delivers the maximum 222 μg/mL dose of fentanyl. Q0 represents the estimated fentanyl intake at zero cost, and α represents demand elasticity. Higher values of α represent higher sensitivity to increasing cost.

### Modeling Estimated Brain-Cocaine Concentration

Whole-brain levels of fentanyl were estimated using a two-compartment model for rats (Pan et al., 1991). Briefly, we used the equation:

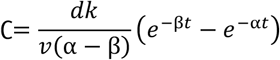

Here, C is the concentration in the brain, d is the dose, k is the rate constant for transfer from the body to the brain (0.233), v is volume of the brain (0.15 l/kg), α and β (0.553 and 0.0671, respectively) are constants representing the flow of fentanyl between the body and brain compartments and the elimination of fentanyl from the body, and t is the time in min since the last infusion. All constants were calculated using methodology derived from intraperitoneal cocaine experiments, but with a larger elimination coefficient (kel=0.175) to account for the relatively longer half-life of fentanyl (Pan et al, 1991).

### Ultrasonic Vocalization Analysis

Mems ultrasonic microphones (M500-384; Pettersson Elektronik) were used to record ultrasonic vocalizations (USVs) during oral fentanyl SA once per training phase per animal. The microphone was inserted into a hole in the SA chamber and was enclosed in a Plexiglas tube surrounded by sound dampening foam. Recording sessions began 1 minute prior to the SA session. Sonorous activity was recorded for 1 hour at a 250-kHz sampling frequency (16 bits) using SASLab Lite software (Avisoft Bioacoustics) and stored for offline analysis. Nearly all USVs occur within the first 30 minutes of a session, so recordings were shorter than SA sessions to save on storage and analysis time. USVs were automatically detected from 18-95 kHz using DeepSqueak v3 (Coffey et al., 2019). All detection files were manually reviewed, and call selections were further refined. USVs were then parameterized and automatically clustered using a combination of variational autoencoder embeddings and contour frequency parameters. This new method incorporates recent advancements in machine vision based latent feature extraction (Goffinet et al., 2021) with traditional contour-based clustering (Coffey et al., 2019) to improve classification. Clusters were chosen by running iterative K-means on the extracted features (K=1 to 100) and using the “ elbow method” to determine an optimal K (K=12). Call clusters were given descriptive labels after unsupervised clustering to aid in discussion of and comparison between call types. Calls were classified as either positive affective (principal freq. > 38 kHz) or negative affective (principal freq. < 38 kHz). Clusters were visualized using UMAP, and differences in call type use were analyzed as a function of session type and session time (McInnes et al., 2020).

### Surgical Methods

For fiber photometry surgeries, rats were anesthetized with 1–3% isoflurane, and operated on using a custom robotic stereotaxic instrument (Coffey et al., 2013). Animals received 500nL injections of pGP-AAV1-syn-jGCaMP7f-WPRE (#104488, Addgene) at 100nL/min into the right Lateral Habenula (LHb; Males: A/P -3.7, M/L 0.95, D/V -5.2; Females: scaled*0.95) using a 33g needle (NF33BV-2, WPI). During the same surgery, a borosilicate optical fiber (MFC_400/430-0.66, Doric Lenses) was inserted directly above the injection site (Males: D/V -4.8; Females: scaled*0.95) and secured to the skull with jeweler’ s screws, cyanoacrylate, and dental acrylic (Figure 5A). After surgeries, rats were given meloxicam (2.5 mg/kg s.c.) for pain management and monitored daily for at least 3 days. Accuracy of injection coordinates was confirmed by visualization of GFP in the LHb. These injection volumes and coordinates were optimized to produce selective transduction of LHb neurons with minimal expression in adjacent regions (Figure 5B).

### Fiber Photometry Methods

Fiber photometry was utilized in the following experiments to record calcium transients and analyze LHb activity during pertinent behavioral events. For all photometry experiments, animals were connected to a turnkey fiber photometry system (RZ10x, TDT) through a fiber optic rotary joint (FRJ_1×1_PT_400-0.57, Doric Lenses), allowing complete freedom of motion for the animals. During behavioral experiments, photometry data was collected using an isosbestic 405 nm LED as well as the signal generating 465 nm LED (Figure 1C). Photometry data (10 ms resolution) from both 465 and 405 nm signals was lightly smoothed (lowess, span=0.01%) and fit to a polynomial curve (n=8) to eliminate drift. The 405 nm control signal was scaled to the 465 nm signal, and the percent change in fluorescence over the 405 control signal was calculated (Figure 5D; %ΔFF=(465-405)/405)*100). This methodology allows for control of motion artifacts and comparison across subjects with variable raw signal voltages. Group change in calcium activity was analyzed be via event related transients (ERT), by averaging %ΔFF around repeated similar events such as a lever press. These analyses are detailed below in the statistics section.

### Histology

To confirm GCaMP7f expression and fiber location after the completion of the experiment, animals were deeply anesthetized with Phenytoin/Pentobarbital and transcardially perfused with ice cold phosphate buffered saline (PBS), followed by 4% paraformaldehyde (PFA). Brains were post-fixed at 4°c for 24h in 4% PFA, followed by 72h in 30% sucrose in PBS. Brains were sectioned at 30μm on a cryostat and mounted to charged slides and stored at room temperature (RT). Immunohistochemistry for GCaMP7f (GFP) was performed directly on slides. Briefly, sections were outline with a hydrophobic barrier (ImmEdge, Vector Labs), and rehydrated in PBS for 5m at RT. Sections were then blocked with 5% bovine serum (BSA) in PBS+0.1% Triton for 1h at RT. Sections were then incubated in a rabbit polyclonal GFP antibody (#A-11122, ThermoFisher) at 1:1000 concentration in the 5% BSA blocking solution for 1h at RT. Sections were then rinsed in PBS 3x for 5m at RT, followed by incubation in secondary antibody (#A-11034, ThermoFisher) at 1:500 concentration for 1h at RT. Sections were then rinsed again in PBS 3x for 5m at RT and finally cover slipped with ProLong Gold + DAPI (#P36931, ThermoFisher). Histological images were acquired at 20x on a widefield microscope (BZ-X800, Keyence) and assessed for expression and fiber placement.

### Statistical Analyses

Statistical analyses for oral fentanyl SA needed to account for repeated measures over time, between groups difference, and nesting within individual animals. To accomplish this, all stats were run using linear mixed-effects modeling via MATLAB’ s fitlme function with the following base formula:

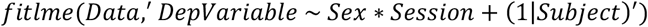

Analysis of variance for each linear mixed-effects model was calculated using the Satterthwaite approximation for degrees of freedom and MATLAB’ s anova() function with the following format:

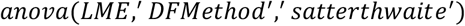

For photometry analysis, it is valuable to identify event related transients (ERTs), and there are numerous ways to accomplish this with varying advantages and disadvantages (Jean-Richard-dit-Bressel et al., 2020). Here we used the bootstrapped confidence intervals (bCI) approach, which is powerful, temporally defined, and non-parametric, but lacks exact p-values. Briefly, to identify ERTs using bCI, average %ΔFF was calculated in the 10s surrounding each event for each subject. 95% CIs were calculated via bootstrapping (randomly resampling with replacement) from the dataset and obtaining a bootstrap estimate from the sampling. This method makes no assumptions about the underlying distribution and has a narrowness bias for small sample sizes, making it relatively conservative. Periods where the bCI does not contain the null (%ΔFF = 0, minimum span = 2s) can be flagged as “ significant”, i.e., indicative of an ERT.

## Acknowledgments

The authors thank Michele Kelly & David Bergkamp for their administrative support and editing. This project was funded by NIDA grant K99DA052571, the UW NAPE Center pilot grant DA048736-03, DA052618-02, and DA052618-01.

## Code and Data Availability

All code and processed data used to generate figures and statistics for this manuscript are available on our [GitHub](https://github.com/DrCoffey/Oral-Fentanyl-Manuscript). A stable release and corresponding DOI will be generated for this data upon acceptance of this manuscript. Large raw photometry and audio files can be requested by contacting the corresponding author.

